# Neural sensitization improves encoding fidelity in the primate retina

**DOI:** 10.1101/482190

**Authors:** Todd R. Appleby, Michael B. Manookin

## Abstract

An animal’s motion through the environment can induce large and frequent fluctuations in light intensity on the retina. These fluctuations pose a major challenge to neural circuits tasked with encoding visual information, as they can cause cells to adapt and lose sensitivity. Here, we report that sensitization, a short-term plasticity mechanism, solves this difficult computational problem by maintaining neuronal sensitivity in the face of these fluctuations. The numerically dominant output pathway in the macaque monkey retina, the midget (parvocellular-projecting) pathway, undergoes sensitization under specific conditions, including simulated eye movements. Sensitization is present in the excitatory synaptic inputs from midget bipolar cells and is mediated by presynaptic disinhibition from wide-field amacrine cells. Direct physiological recordings and a computational model indicate that sensitization in the midget pathway supports accurate sensory encoding and prevents a loss of responsiveness during dynamic visual processing.

## INTRODUCTION

The fundamental constraints on sensory coding require that neural circuits adjust their outputs based on the statistical properties of their recent inputs (Srinivasan et al., 1982; Barlow, 1961; Laughlin, 1981). Neurons respond to dynamic inputs using two distinct strategies—adaptation and sensitization. Adapting cells respond to strong stimulation by decreasing their sensitivity and this decrease in responsiveness can persist for several seconds after the stimulus intensity decreases (Baccus and Meister, 2002; Carandini and Ferster, 1997; Kim and Rieke, 2001; Laughlin, 1981; Manookin and Demb, 2006; Smirnakis et al., 1997; Solomon et al., 2004). Thus, adapting cells are relatively insensitive to weak stimuli occurring during these transition periods. Sensitizing cells show the opposite pattern—increasing their responsiveness at these transitions (Kastner and Baccus, 2011; Kastner and Baccus, 2013; Nikolaev et al., 2013). For this reason, adaptation and sensitization are commonly thought to constitute opposing and complementary forms of short-term neural plasticity (Kastner and Baccus, 2011; Kastner and Baccus, 2013).

This hypothesis requires that a sensitizing cell type have an adapting counterpart that encodes common information (Kastner and Baccus, 2011). However, this constraint 36 could potentially decrease the amount of information that can be encoded in an neural ensemble and increase the metabolic demands on a sensory tissue (Laughlin, 1981; Balasubramanian et al., 2001). Alternatively, adaptation and sensitization could be signatures of fundamentally distinct neural coding strategies (Młynarski and Hermundstad, 2018). Further, these alternative hypotheses are not mutually exclusive—adapting and sensitizing cells could mirror each other in some species and neural pathways and not in others, depending on the particular coding and metabolic constraints in those systems (Laughlin, 1981; Barlow, 1961; Levy and Baxter, 1996; Balasubramanian et al., 2001). However, given that neural sensitization was only recently discovered, relatively little is known about its roles in neural information processing.

To address this issue, we recorded from five types of output neurons in the macaque monkey retina—broad thorny, On and Off parasol (magnocellular-projecting), and On and Off midget (parvocellular-projecting) ganglion cells. These cells have well described roles in visual processing and no known functional counterparts. We studied how these cells responded to global fluctuations in contrast and other stimulus statistics. We report that whereas broad thorny and parasol cells strongly adapted, midget cells sensitized—increasing their responsiveness to certain types of visual stimulation, including high contrast and simulated eye movements. Synaptic current recordings revealed that this increased sensitivity was present in the excitatory input from midget bipolar cells and was mediated by presynaptic disinhibition. A computational model based on synaptic input recordings further indicated that this increase in sensitivity greatly enhanced the fidelity of encoding natural scenes. Moreover, the lack of an adapting counterpart to midget cells indicates that sensitizing circuits perform a distinct role in primate retina relative to that observed in other vertebrate neural systems (Kastner and Baccus, 2011; Kastner and Baccus, 2013; Nikolaev et al., 2013; Cohen-Kashi Malina et al., 2013).

## RESULTS

The midget pathway of the primate retina is commonly believed to lack short-term plasticity mechanisms such as contrast gain control. This belief is based on reports that midget cells did not exhibit noticeable changes in responsiveness following transitions from high to low contrast regimes (Solomon et al., 2004; Benardete et al., 1992). The assay used to measure adaptation was a sinusoidally modulated drifting grating with bar widths tuned to the size of the midget cell receptive field center, which is narrower than many other retinal cell types. Thus, if plasticity in the midget pathway depended on mechanisms with broader spatial tuning, this assay would not engage such mechanisms.

To determine whether short-term plasticity in the midget pathway depended on the spatial properties of the stimulus, we repeated this assay while varying the spatial tuning of the gratings. At the offset of high contrast, midget cells did not exhibit a notable change in firing relative to the period that preceded high-contrast stimulation (Solomon et al., 2004; Benardete et al., 1992) (Figure 1C; spatial frequency, 3.5 cycles degree^−1^). To determine whether this lack of either adaptation or sensitization persisted across a range of stimulus conditions, we varied the spatial frequency content of the drifting gratings. Following the offset of low spatial frequency gratings, most midget cells showed an increase in spiking relative to the period preceding grating onset (Figure 1D; spatial frequency, 0.35 cycles degree^−1^). This increase in spiking following high contrast is characteristic of the contrast sensitization observed in other vertebrate retinas (Kastner and Baccus, 2011; Kastner and Baccus, 2013; Nikolaev et al., 2013). The presence of sensitization at low spatial frequencies suggested that sensitization depended on the ability to engage elements in the midget cell receptive field with broad spatial tuning relative to the midget bipolar cell.

**Figure 1.**
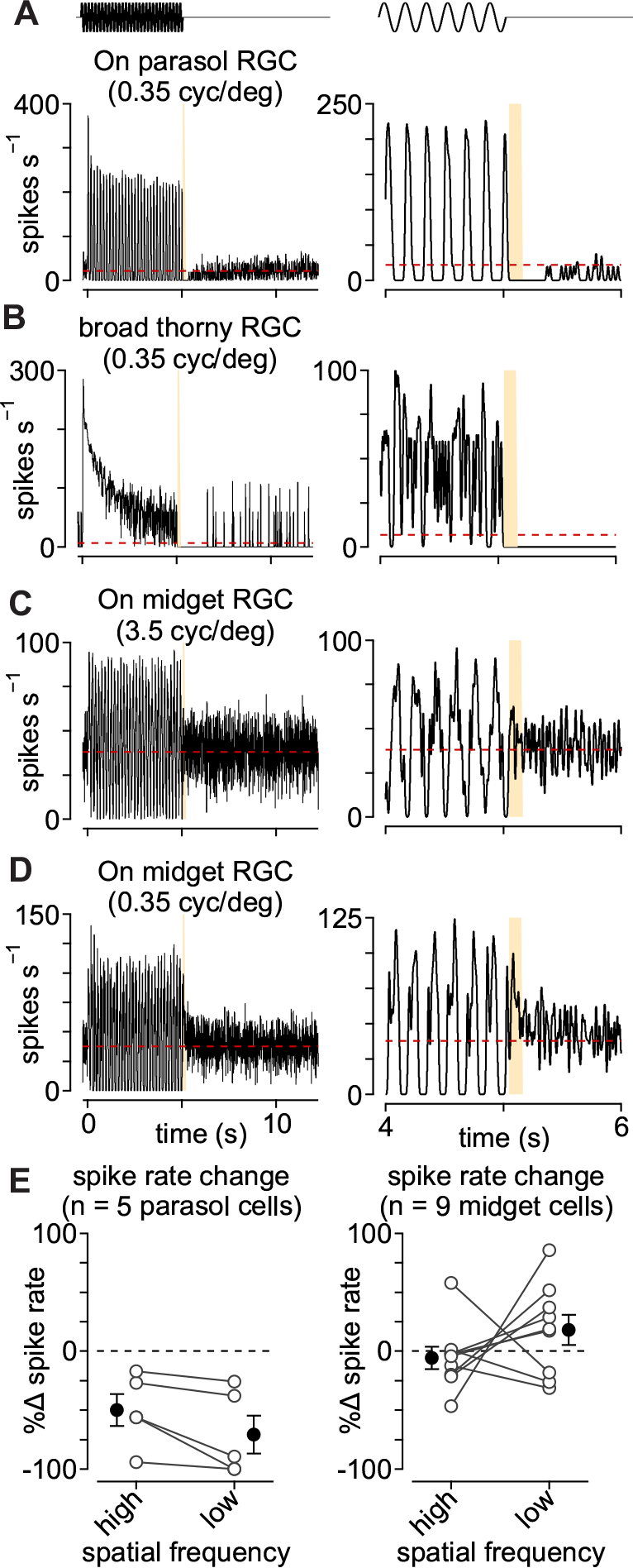
Parasol and midget cells exhibit opposing forms of plasticity. (A) Spike rate in an On parasol ganglion cell to a low spatial frequency drifting grating presented for five seconds (temporal frequency, 6 Hz; spatial frequency, 0.35 cycles degree^−1^). After the offset of high contrast, the spike rate declined below the level prior to grating onset (red dashed line). *Right*, zoom of transition period. (B) Same as (A) in a broad thorny (On-Off type) ganglion cell. (C) Same as (A) in an On midget ganglion cell to a high spatial frequency grating (3.5 cycles degree^−1^). (D) Spike responses from the same cell as in (C) to a low spatial frequency grating (0.35 cycles degree^−1^). (E) Change in spike rate for the period directly after grating offset relative to period prior to grating onset in parasol (left) and midget ganglion cells (right).

Parasol and broad thorny cells responded very differently than midgets. At the transition from high to low contrast, these cells showed a pronounced decrease in spiking relative to the period before the grating turned on and several seconds were required for the spike rate to recover (Figure 1A, B; high contrast, 1.0; low contrast, 0.0; spatial frequency, 0.18-3.5 cycles degree^−1;^ grating size, 730 μm × 730 μm). This behavior is characteristic of contrast adaptation—during periods of high contrast, circuit mechanisms reduce the gain to avoid saturation and the gain remains low for several seconds following the transition to a low-contrast regime (Chander and Chichilnisky, 2001; Benardete and Kaplan, 1999; Solomon et al., 2004).

### Wide-field stimulation evokes contrast sensitization in midget ganglion cells

Our next goal was to determine how this putative wide-field component of the midget cell receptive field contributed to contrast coding. To accomplish this goal, we sought a more spatially and temporally precise assay of sensitivity following wide-field adaptation. Contrast tuning of parasol and midget cells was determined with spots centered on the receptive field (duration, 0.1 s; parasol diameter, 80-200 μm; midget diameter, 40-80 μm). Contrast responses were measured in isolation (unadapted condition) or 50-100 ms following the offset of an adapting stimulus (adapted condition). The adapting stimulus was a large, high-contrast spot modulated at 20-30 Hz (diameter, 730 μm; contrast, 0.5-1.0, duration, 1.25 s). Presentations of the adapted and unadapted stimuli were interleaved to account for any potential variability in cellular responses over time.

Example spike responses to this stimulus paradigm are shown in Figure 2. Parasol cells increased their spike rate at the onset of the adapting stimulus and the spike rate quickly decreased to a steady-state rate by ~0.25 s. Test flashes presented after the offset of the adapting stimulus evoked fewer spikes relative to the unadapted control (Figure 2A). Both of these patterns—a transient increase in spike rate following the transition to high contrast and a decrease in spiking after the transition to low contrast—are characteristic of cells undergoing contrast adaptation (Kim and Rieke, 2001; Baccus and Meister, 2002; Brown and Masland, 2001).

**Figure 2.**
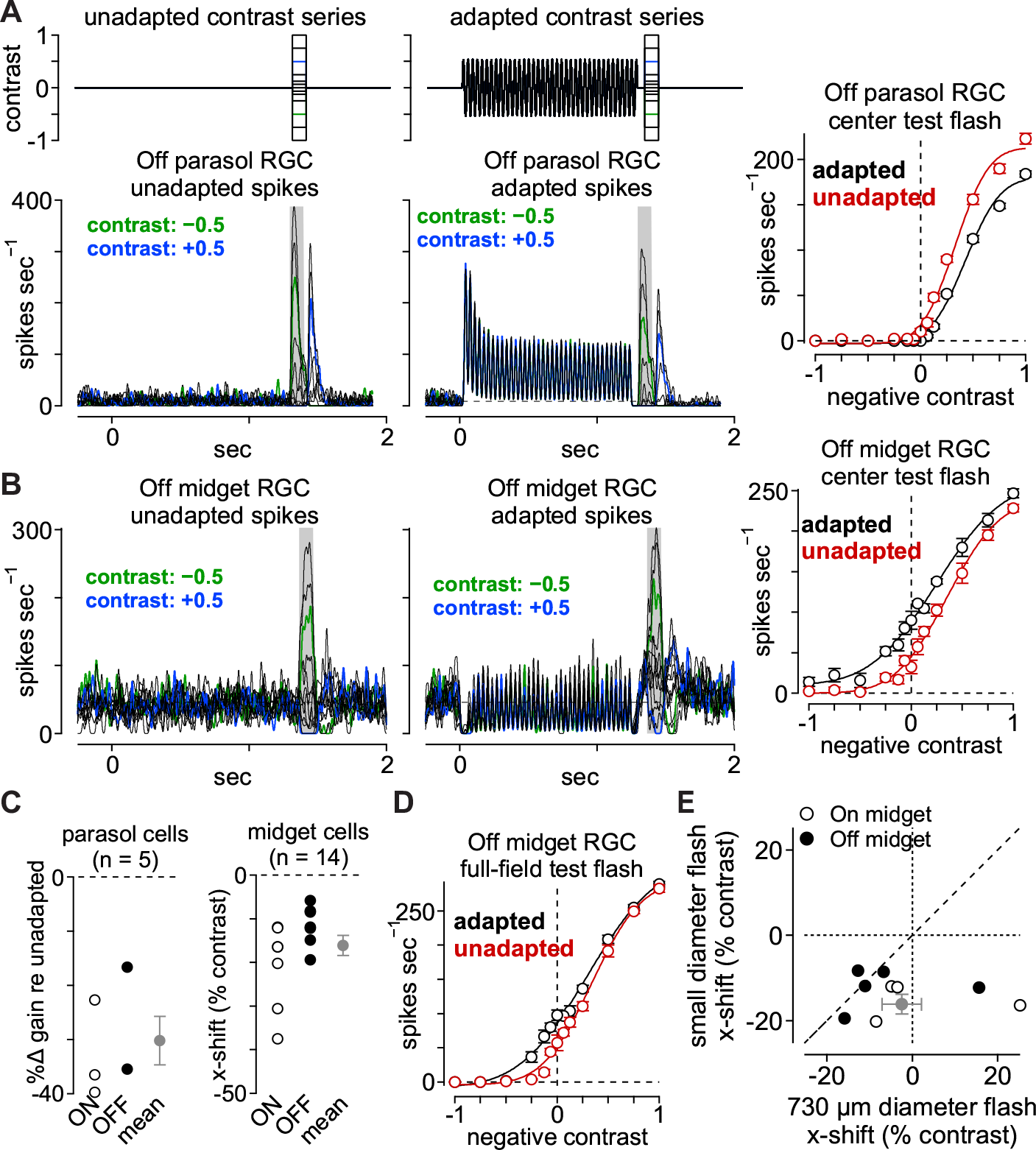
Midget ganglion cells display contrast sensitization. (A) Spike responses from an Off parasol ganglion cell to a series of spots centered over the receptive-field. Spots were either presented alone (left) or 50 ms following the offset of an adapting stimulus (right). Shaded regions indicate sampling windows. *Right*, Average spike rate across the shaded regions. The wide-field adaptation evoked a decrease in the slope (gain) of the contrast-response curve (black) relative to the unadapted control condition (red). (B) Same as (A) for an Off midget ganglion cell. *Right*, Average spike rate across the shaded regions. The wide-field adaptation evoked a leftward shift in the contrast-response curve (black) relative to the unadapted control condition (red). (C) *Left*, Population data showing the change in slope (gain) for the adapted condition relative to the unadapted condition in On (open circles) and Off (closed circles) parasol ganglion cells (n = 5). *Right*, Population data showing the *x*-axis shift for adapted relative to unadapted conditions for small-diameter test flashes in On (open circles) and Off (closed circles) midget cells (n = 14). Gray circle and bars indicate mean ± SEM. (D) Average spike rate evoked by wide-field test flashes for the Off midget cell in (B). (E) Population data showing the *x*-axis shift for adapted relative to unadapted conditions for wide-field test flashes versus small-diameter test flashes in On (open circles) and Off (closed circles) midget cells. Gray circle and bars indicate mean ± SEM.

We modeled the variation in the contrast-response function following the adapting stimulus as a change in the slope (gain) and a horizontal shift relative to the control condition (see Methods). Following the adapting stimulus, parasol cells showed a large decrease in gain (−30.2 ± 4.5%; n = 5 cells; p = 1.3 × 10^−3^; Wilcoxon signed rank test, here and below) and a small rightward horizontal shift (+3.6 ± 1.5% contrast; p = 3.0 × 10^−2^) relative to the unadapted control (Figure 2C). This result confirms previous reports that parasol cells readily adapt to contrast by continuously adjusting their sensitivity to match the statistics of incoming visual inputs (Chander and Chichilnisky, 2001; Solomon et al., 2004; Benardete et al., 1992).

Midget cells showed several striking differences relative to the pattern observed in parasol cells. First, the decrease in gain was much smaller in midget cells (−5.4 ± 4.3%; n = 14 cells; p = 0.12). Second, an increase in spike rate was observed at the offset of the adapting stimulus relative to the unadapted control (Figure 2B). This increase in spiking was evident at all contrasts tested including the zero-contrast condition in which the spot intensity was equal to the average background intensity. The elevation in spiking following the adapting stimulus produced a leftward shift in the contrast-response curve relative to control (−16.2 ± 2.3% contrast; p = 5.2 × 10^−6^). A negative horizontal shift value occurred when the adapted curve was shifted to the left of the control curve and this indicated that a weaker stimulus was required to elicit the same spike response from a midget cell following the adapting stimulus. This observation was consistent with previous reports demonstrating that a decreased spike threshold, increased baseline response, and slight decrease in gain are characteristic of contrast sensitization (Kastner and Baccus, 2011; Kastner and Baccus, 2013).

### Contrast sensitization is reduced for wide-field stimulation

Midget cells show narrow receptive-field centers with strong input from the receptivefield surround (Crook et al., 2011; De Monasterio and Gouras, 1975; Derrington et al., 1984). Thus, the effect of sensitization may be diminished following the adapting stimulus depending on the relative influences of the direct midget bipolar cell input and wide-field mechanisms in contrast sensitization. To determine whether contrast sensitization varied with the size of the test flash, we repeated the adaptation experiment but used wide-field test flashes to measure the contrast tuning of midget cells (diameter, 730 μm). The wide-field test flash evoked a slight leftward shift for the adapted condition relative to control, but this shift was much smaller than was observed for the small-diameter test flash in the same cell (compare Figure 2B, D). This trend held true across midget cells—horizontal shifts were more negative for the small-diameter test flash than for the wide-field test flash in the same cell and these shifts were not statistically significant for the wide-field test flashes (x-shift, −2.4 ± 4.6% contrast; p = 0.30; gain change, −5.8 ± 5.8%; n = 9 cells; p = 0.17; Figure 2E). These data indicated that the relative activations of narrow-field and wide-field mechanisms during and following the adapting stimulus were critical to contrast sensitization in midget ganglion cells. Moreover, this result agrees well with previous findings that did not report contrast sensitization to wide-field noise (Chander and Chichilnisky, 2001).

### Time course for the onset and persistence of sensitization

We next sought to determine the amount of stimulation needed to evoke sensitization and also how long sensitization persisted after its onset. To determine the stimulation period needed to initiate sensitization, we varied the presentation time of the adapting stimulus and measured the change in spike rate relative to the unadapted control (adaptation duration, 0.25-1.25 s; contrast, ±0.5; duration, 0.1 s). For each period of adaptation, midget cells showed an elevation in spiking relative to unadapted controls (Figure 3A). Thus, sensitization could be elicited even with fairly brief stimulus presentations.

**Figure 3.**
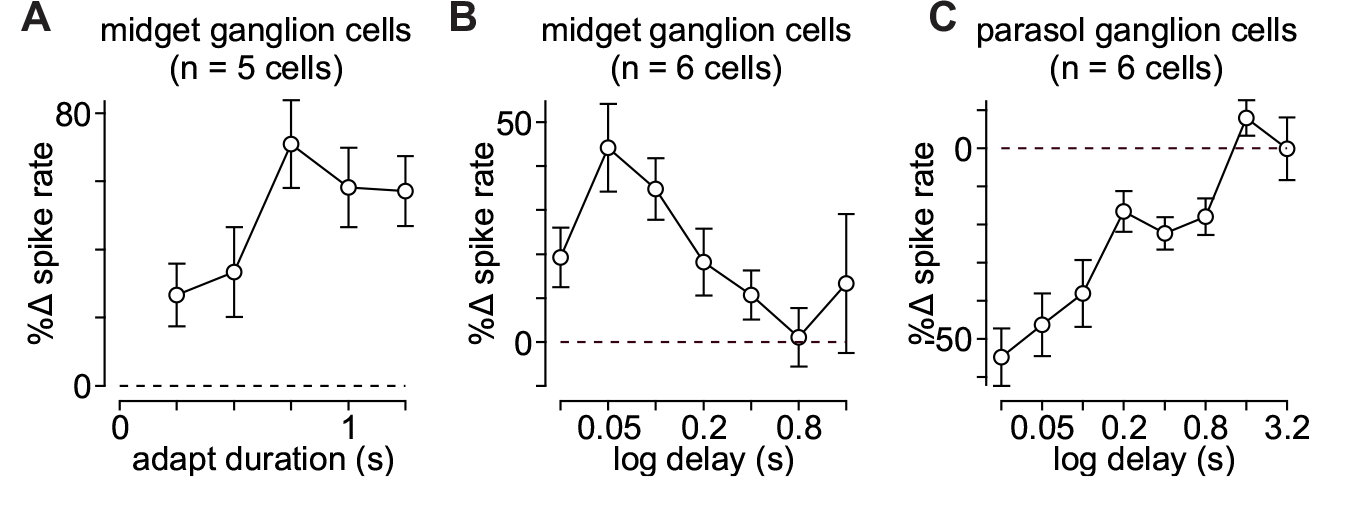
Time course of contrast sensitization and adaptation. (A) Change in spike rate for the adapted condition relative to unadapted control for adaptation periods (contrast, ±0.25-0.5; delay 0.05 s). Adaptation period was varied between 0.25-1.25 s (*x*-axis). (B) Duration of contrast sensitization in midget ganglion cells. Test flashes (contrast, ±0.25-0.5) were presented at different delays (*x*-axis) following the offset of an adapting stimulus. Percent change in spike rate for the adapted condition relative to the unadapted condition is shown on the *y*-axis. (C) Same as (B) for parasol ganglion cells. Error bars indicate mean ± SEM.

To determine the time course of sensitization in midget cells, we measured spot responses at different times following the offset of the adapting stimulus (delay, 0.025-1.6 sec; contrast, ±0.5; duration, 0.1 sec). Relative to the unadapted control, the adapting stimulus elicited higher spike rates to the test flash in midget cells at delays of 0.025 seconds (Figure 3B). This elevation in spiking, characteristic of sensitization, was greatest 0.05-0.1 seconds after the offset of the adapting stimulus. Parasol cells, on the other hand, showed a reduction in spiking to the same stimulus that persisted for approximately one second (Figure 3C). Together, these data indicated that sensitization in midget cells could be elicited even with fairly brief stimulus presentations and that it persisted for several hundred milliseconds.

### Sensitization enhances chromatic processing in midget cells

Midget ganglion cells in the central retina exhibit strong chromatic opponency which is formed from differential input from long-wavelength cones (L cones) and middle wavelength cones (M cones) to the receptive-field center and surround (Crook et al., 2011; De Monasterio and Gouras, 1975; Derrington et al., 1984). To determine whether sensitization affected chromatic processing, we measured contrast responses in midget cells with purely chromatic (isoluminant) test flashes following the adapting stimulus.

Isoluminant (equiluminant) stimuli are commonly employed to study color mechanisms in isolation. These stimuli are created by modulating L and M cones in opposing phases to silence achromatic mechanisms that sum inputs from these cone types (i.e., L+M). We measured contrast-responses to purely chromatic (isoluminant) flashes (duration, 0.1 sec) in the presence or absence of an achromatic adapting stimulus, as above. As with the achromatic stimuli, the adapting stimulus elicited a leftward shift to chromatic test contrasts (Figure 4). This shift was reminiscent of that observed for achromatic stimulation (−11.3 ± 4.1% contrast; n = 8 cells; p = 1.5 × 10^−2^). These data indicated that contrast sensitization enhanced both achromatic and chromatic processing in midget cells.

**Figure 4.**
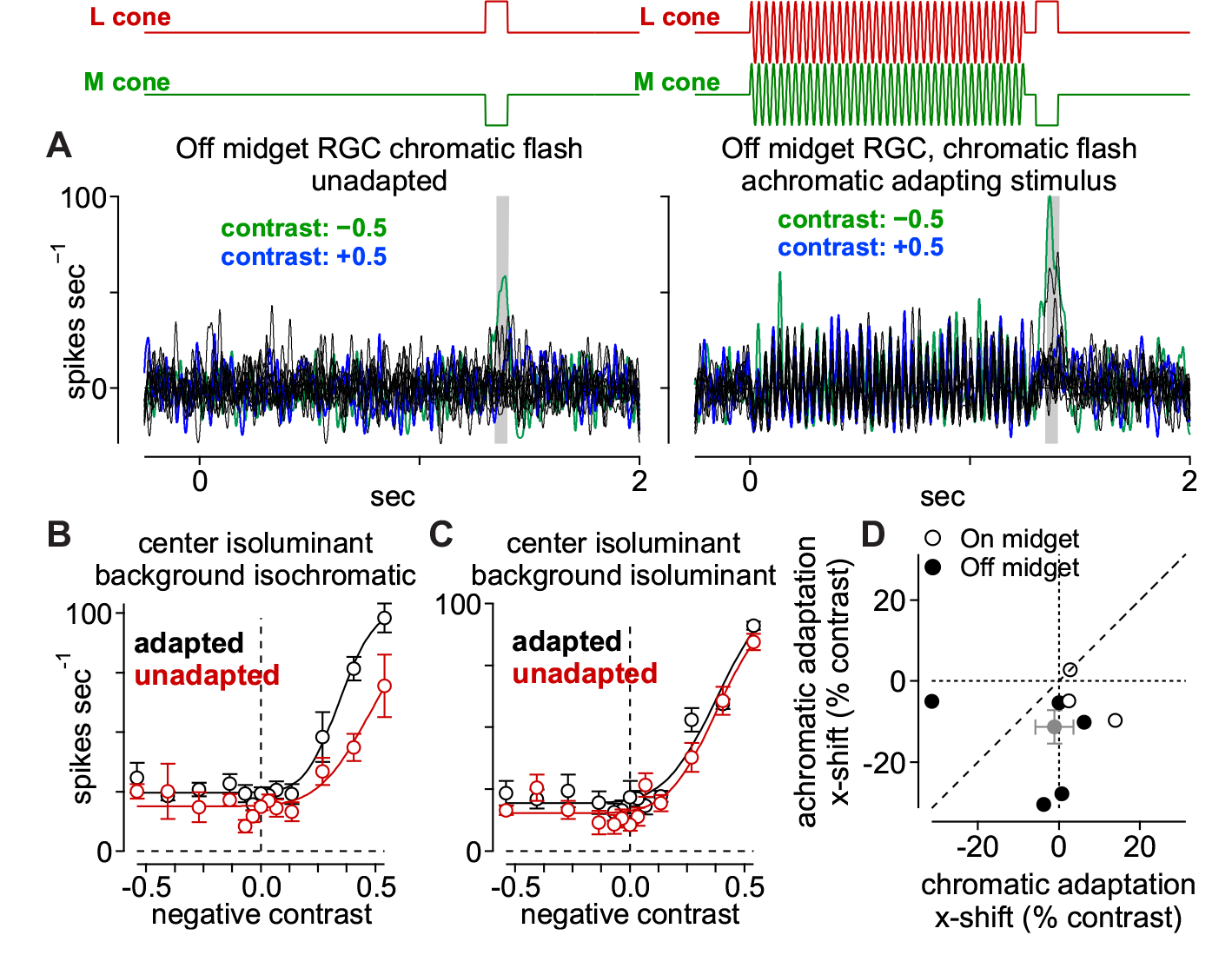
Sensitization arises from an achromatic mechanism. (A) Spike responses from an Off midget ganglion cell to a chromatic (isoluminant) contrast series. Spots were either presented alone (left) or 50 ms following the offset of an achromatic adapting stimulus (right). Shaded regions indicate sampling windows. (B) Average spike rate across the shaded regions indicated in (A). Achromatic adaptation evoked a leftward shift in the contrast-response curve (black) relative to the unadapted control condition (red) for the chromatic test flash. (C) Same as (B) for a chromatic adapting stimulus. The chromatic adapting stimulus did not evoke change in the contrast-response curve relative to control. (D) Population data showing the *x*-axis shift for adapted relative to unadapted conditions for a chromatic adapting stimulus (*x*-axis) relative to an achromatic adapting stimulus (*y*-axis) in On (open circles) and Off (closed circles) midget cells. Gray circle and bars indicate mean ± SEM.

While chromatic processing was affected by sensitization, the observation that an achromatic adapting stimulus was sufficient to evoke sensitization indicated that chromatic circuits were not necessary to elicit the phenomenon. These data did not, however, rule out contributions from purely chromatic mechanisms to contrast sensitization.

### Sensitization does not arise from a chromatic mechanism

To determine whether such a chromatic mechanism contributed to the observed contrast sensitization, we presented a chromatic adapting stimulus. This stimulus was specifically designed to modulate chromatic mechanisms that differentiate L- and M-cone inputs (L−M; isoluminant) while silencing achromatic mechanisms that sum inputs from the L- and M-cone pathways (L+M; isochromatic). Following the adapting stimulus, an isoluminant contrast series was used to measure the input-output relationship. In the same cell, we compared chromatic contrast-responses following a chromatic (L−M) or achromatic (L+M) adapting stimulus.

The achromatic adapting stimulus produced a leftward shift in the chromatic contrast-response relation. The chromatic adapting stimulus, however, produced no such shift (x-shift, −1.1 ± 4.7% contrast; n = 8 cells; p = 0.41). We interpreted this result as evidence that contrast sensitization arose from an achromatic mechanism in the midget cell receptive-field. Moreover, given the role of horizontal cells in forming the L-versus-M opponent receptive-field surround, these data excluded horizontal cells as the source of sensitization in the midget pathway (Crook et al., 2011).

### Sensitization is present in excitatory synaptic input from midget bipolar cells

The experiments above found contrast sensitization in the spike output of midget ganglion cells. Our next goal was to understand the circuit mechanisms mediating sensitization. To accomplish this goal, we measured the direct excitatory and inhibitory synaptic inputs to midget ganglion cells with whole-cell, voltage-clamp recordings (see Methods). Excitatory currents were isolated by holding a cell’s membrane voltage at the reversal potential for inhibition (−70 mV), and likewise, inhibitory currents were recorded at the excitatory reversal potential (0 mV). An increase in excitatory input to a cell was indicated by a more negative (inward) current relative to the leak current. Indeed, the adapting stimulus evoked larger inward excitatory currents relative to the unadapted control at all contrasts tested (Figure 5A). Plotting excitatory charge as a function of contrast revealed a similar pattern to that observed in the spike recordings—the adapting stimulus evoked a leftward shift in the contrast-response curve relative to the unadapted control (Figure 5B). On average, the adapting stimulus elicited a horizontal shift of −11% contrast (−11.3 ± 4.2% contrast; n = 8 cells; p = 1.95 × 10^−2^). These results indicated that contrast sensitization was present in the excitatory synaptic input from midget bipolar cells to midget ganglion cells.

**Figure 5.**
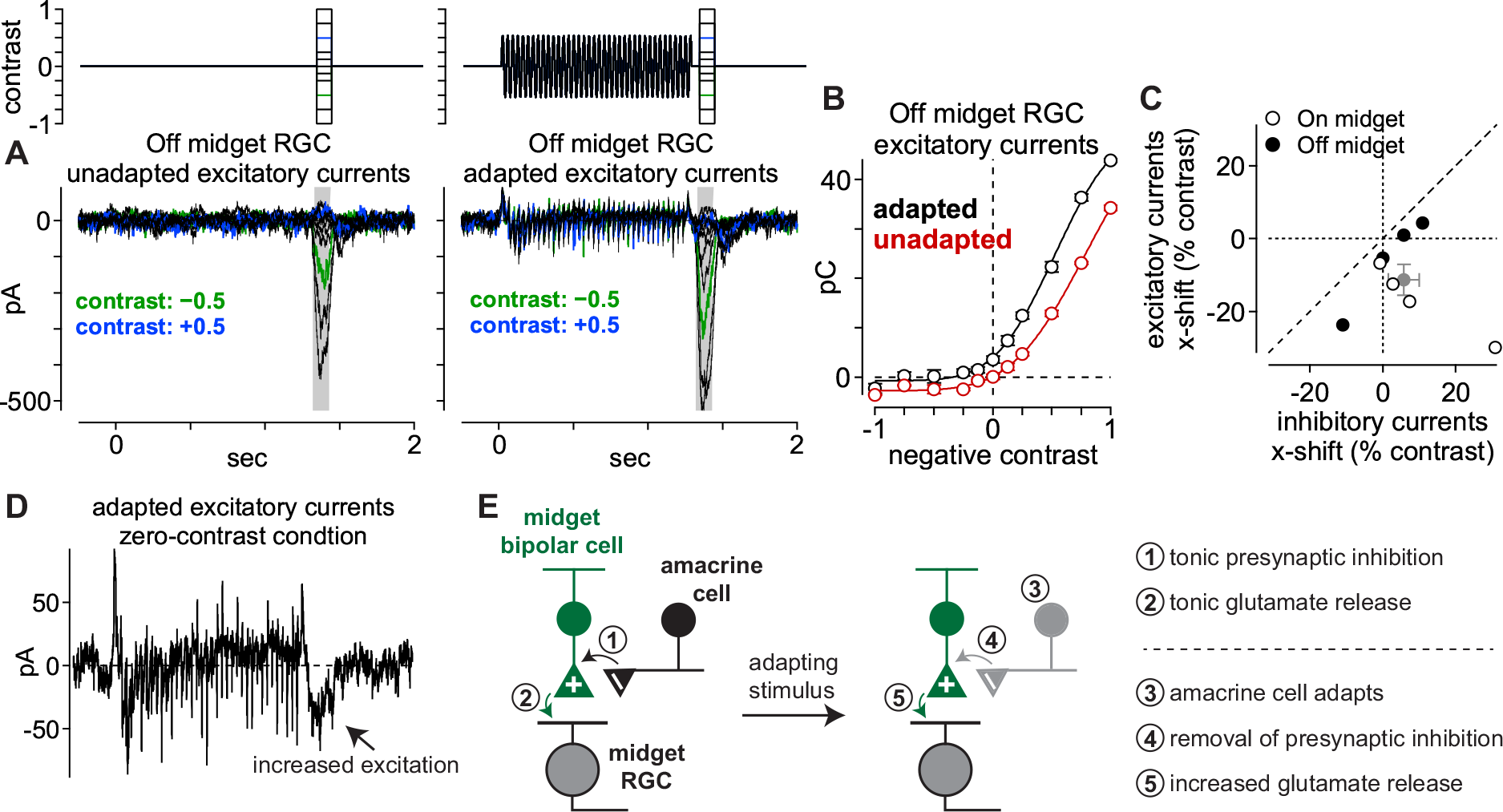
Sensitization present in excitatory synaptic input from midget bipolar cells. (A) Excitatory currents from an Off midget ganglion cell to a series of spots (diameter, 40-80 μm) centered over the receptive field. Spots were either presented alone (*left*) or 50 ms following the offset of an adapting stimulus (*right*; diameter, 730 μm). Shaded regions indicate sampling windows. (B) Average spike rate across the shaded regions indicated in (A). The wide-field adaptation evoked a leftward shift in the contrast-response curve (black) relative to the unadapted control condition (red). (C) Population data showing the x-axis shift for adapted relative to unadapted conditions for excitatory versus inhibitory synaptic currents in On (open circles) and Off (closed circles) midget cells. Mean values are shown in gray. Error bars indicate mean ± SEM. (D) Excitatory current recordings from the Off midget cell in (A) under the condition in which the stimulus intensity returned to the mean luminance after the offset of the adapting stimulus and an additional test flash was not presented (zero-contrast condition). A sustained increase in excitatory current was observed at the offset of that stimulus. (E) Proposed model for contrast sensitization in midget bipolar cells.

We also tested for the presence of sensitization in the inhibitory synaptic inputs to midget cells. Unlike the pattern observed in spiking and excitatory currents, the adapting stimulus did not consistently elicit leftward shifts in the inhibitory contrast response functions relative to control (+5.8 ± 4.3% contrast; n = 8 cells; p = 0.25; Figure 5C). These data indicated that contrast sensitization arose at or prior to the level of glutamate release from midget bipolar cells. This finding was consistent with the circuit model for contrast sensitization in bipolar cells in the retinas of fish, salamander, mice, and rabbits (Kastner and Baccus, 2013; Kastner and Baccus, 2011; Nikolaev et al., 2013). This model posited a mechanism in which a strongly adapting amacrine cell drove sensitization by a mechanism of presynaptic inhibition at the bipolar cell terminal (Kastner and Baccus, 2013). During the adapting stimulus, the amacrine cell adapted such that it decreased release of inhibitory neurotransmitter to the bipolar cell synaptic terminal relative to the tonic level following stimulus offset. This presynaptic disinhibition, in turn, depolarized the bipolar cell synaptic terminal, allowing the cell to utilize its full dynamic range in signaling via glutamate release to postsynaptic ganglion cells.

Cleanly measuring the effects of presynaptic inhibition on circuit function has proven exceedingly difficult as use of inhibitory receptor antagonists typically cause many offtarget effects that make data interpretation highly tenuous (Cook et al., 1998). Indeed, adding inhibitory antagonists in primate retina evoked significant increases in tonic glutamate release from bipolar cells and changed the contrast polarity of On parasol cells (Manookin et al., 2018). Nonetheless, our spike and whole-cell recordings strongly supported the proposed model in which contrast sensitization arose from disinhibition at the presynaptic bipolar cell terminal (Kastner and Baccus, 2013). First, the lack of sensitization to a purely chromatic (isoluminant) adapting stimulus indicated that sensitization did not arise in the outer retina at the level of horizontal cell feedback (Figure 4). Second, the effect of presynaptic disinhibition was seen in our excitatory current recordings (Figure 5D). In one of our stimulus conditions the test flash contrast was zero such that the stimulus intensity returned to the average background intensity at the offset of the adapting stimulus. Although this stimulus lacked a change in contrast following the adapting stimulus, we observed an increase in excitatory synaptic input (Figure 5D). This response pattern was consistent with a decrease in presynaptic inhibition following the offset of the adapting stimulus, resulting in an increase in glutamate release from midget bipolar cells. Thus, our recordings in midget pathway of primate retina were consistent with the circuit motif proposed in other vertebrate species (Figure 5E; (Kastner and Baccus, 2013)).

### A contrast sensitization model reproduces midget cell responses

Having established the presence of contrast sensitization in midget bipolar cells, we next sought to understand the relevance of this neural computation to visual processing in primates. To accomplish this goal, we developed a computational model of the proposed circuit in which bipolar cell glutamate release was modulated through presynaptic amacrine cell inhibition (Kastner and Baccus, 2013; Kastner and Baccus, 2011; Nikolaev et al., 2013). Model parameters were determined by recording excitatory and inhibitory synaptic current responses from midget ganglion cells to a Gaussian white noise stimulus (see Methods).

We modeled the midget bipolar and presynaptic amacrine cell pathways using the classical linear-nonlinear model with two modifications: 1) adaptation occurred at the amacrine cell output and 2) the amacrine cell output was applied to the bipolar cell model prior to the bipolar cell output nonlinearity (Figure 6A). The model parameters controlling presynaptic sensitization were fit from direct excitatory current recordings. In the same cell from which these parameters were determined, we measured excitatory current responses to the wide-field adapting stimulus (see Figure 5), and the model qualitatively reproduced the increase in excitatory currents following the offset of this adapting stimulus (Figure 6C).

We further tested the model using the drifting grating stimuli presented in Figure 1. The model produced distinct outputs for the high and low spatial frequency gratings. The high frequency grating produced a relatively small response and, as a result, little adaptation in the presynaptic amacrine cell (Figure 6D, middle row). This was due to the broad receptive field center size of the amacrine cell relative to the bars of the grating. Low frequency gratings, however, strongly modulated the amacrine cell and produced significant adaptation; this adaptation, in turn, caused a removal of inhibition at the level of the bipolar cell following the offset of the grating, resulting in sensitization (Figure 6D, bottom row). The model predictions were qualitatively similar to our direct recordings from midget cells, indicating that contrast sensitization in primate retina can be well explained via presynaptic disinhibition as in other species (Kastner and Baccus, 2013; Kastner and Baccus, 2011; Nikolaev et al., 2013).

**Figure 6.**
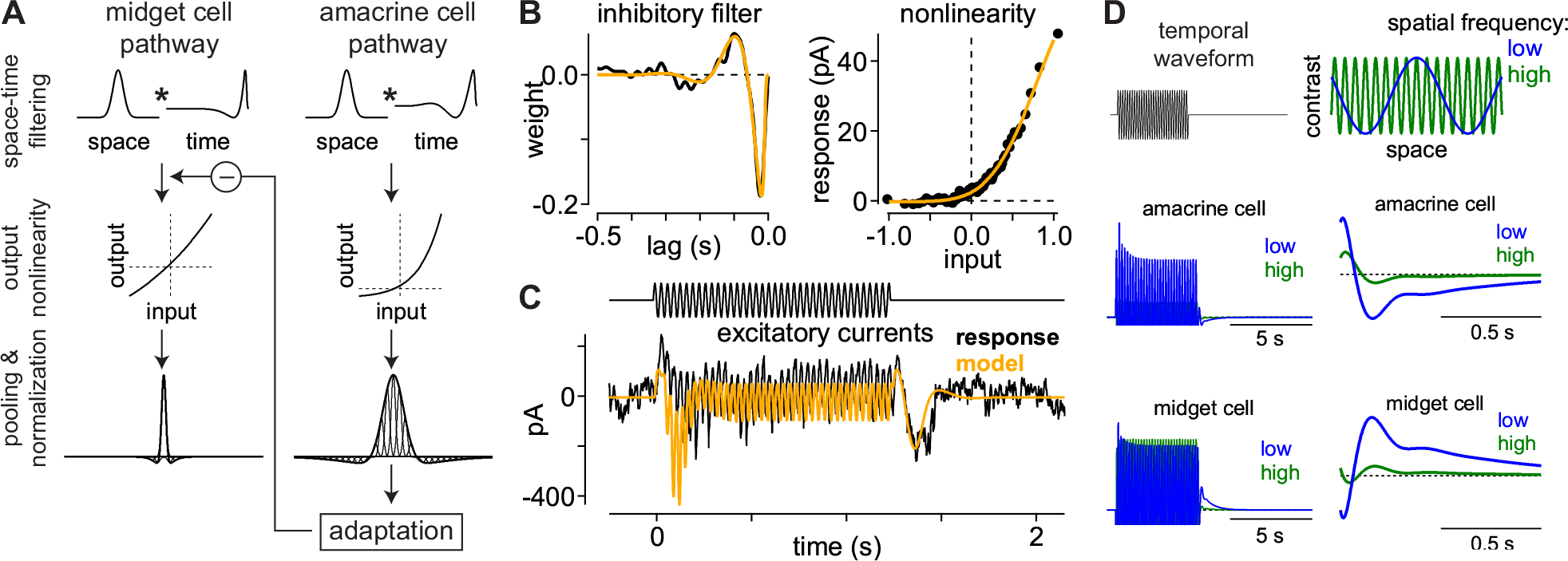
Sensitization model reproduces experimental results. (A) Sensitization model structure. Visual inputs were convolved with a spatiotemporal linear filter comprised of a Gaussian in space and a biphasic filter in time. Signals in the amacrine cell pathway were then passed through an output nonlinearity before passing to the adaptation stage of the model. The output of the amacrine cell model provided inhibitory input to the midget bipolar cell pathway upstream of the bipolar cell output nonlinearity. (B) Inhibitory temporal filter (*left*) and input-output nonlinearity (*right*) determined from noise recordings. These filters were then used as components of the computational model (A). (C) Excitatory current recording from an Off midget ganglion cell to the wide-field adapting stimulus (see Figure 5). Model prediction (orange) was generated from excitatory synaptic current recordings to the noise stimulus in the same cell. (D) Model output for drifting grating stimuli at high and low spatial frequencies.

### Sensitizing circuits more accurately reconstruct natural stimuli than adapting circuits

We next sought to understand how these differing strategies of adaptation and sensitization impacted encoding during naturalistic vision. This was done by testing the ability of adapting and sensitizing models to accurately encode natural scenes. We specifically wanted to determine how accurately downstream visual circuits could reconstruct naturalistic input stimuli based on the outputs of populations of model On and Off midget ganglion cells. The naturalistic stimuli used in the model were taken from the DOVES database—a dataset of eye movements in humans recorded while observing natural images (Van Der Linde et al., 2009). Reconstruction accuracy was determined by calculating the correlation between the stimulus and response of each model (see Methods). Periods of fixation between ballistic eye movements are critically important to visual coding in primates; thus, model performance was separately calculated for the complete movie or for periods of fixation only.

We considered two different decoding models for estimating the stimulus contrast based on the outputs of On and Off midget ganglion cells. The first model utilized a linear decoding scheme in which stimulus contrast was estimated by taking the scaled difference between the On and Off cell outputs. We also tested a quadratic decoding model that squared the On and Off outputs prior to differencing (see Methods). Using these decoders, we compared the performance of the sensitization model with a model in which the midget bipolar underwent contrast adaptation. Regardless of the decoding scheme used, the sensitizing model showed higher accuracy for reconstructing the entire stimulus trajectory than the adapting model (linear *r*^2^: sensitization, 0.81 ± 0.05; adaptation, 0.23 ± 0.07; p = 2.7 × 10^−54;^ quadratic *r*^2^: sensitization, 0.84 ± 0.05; adaptation, 0.45 ± 0.09; p = 2.9 × 10^−54;^ n = 161 movies; mean ± SD; Figure 7C). The sensitizing model also outperformed the adapting model when the analysis was restricted to periods of fixation (Figure 7D).

**Figure 7.**
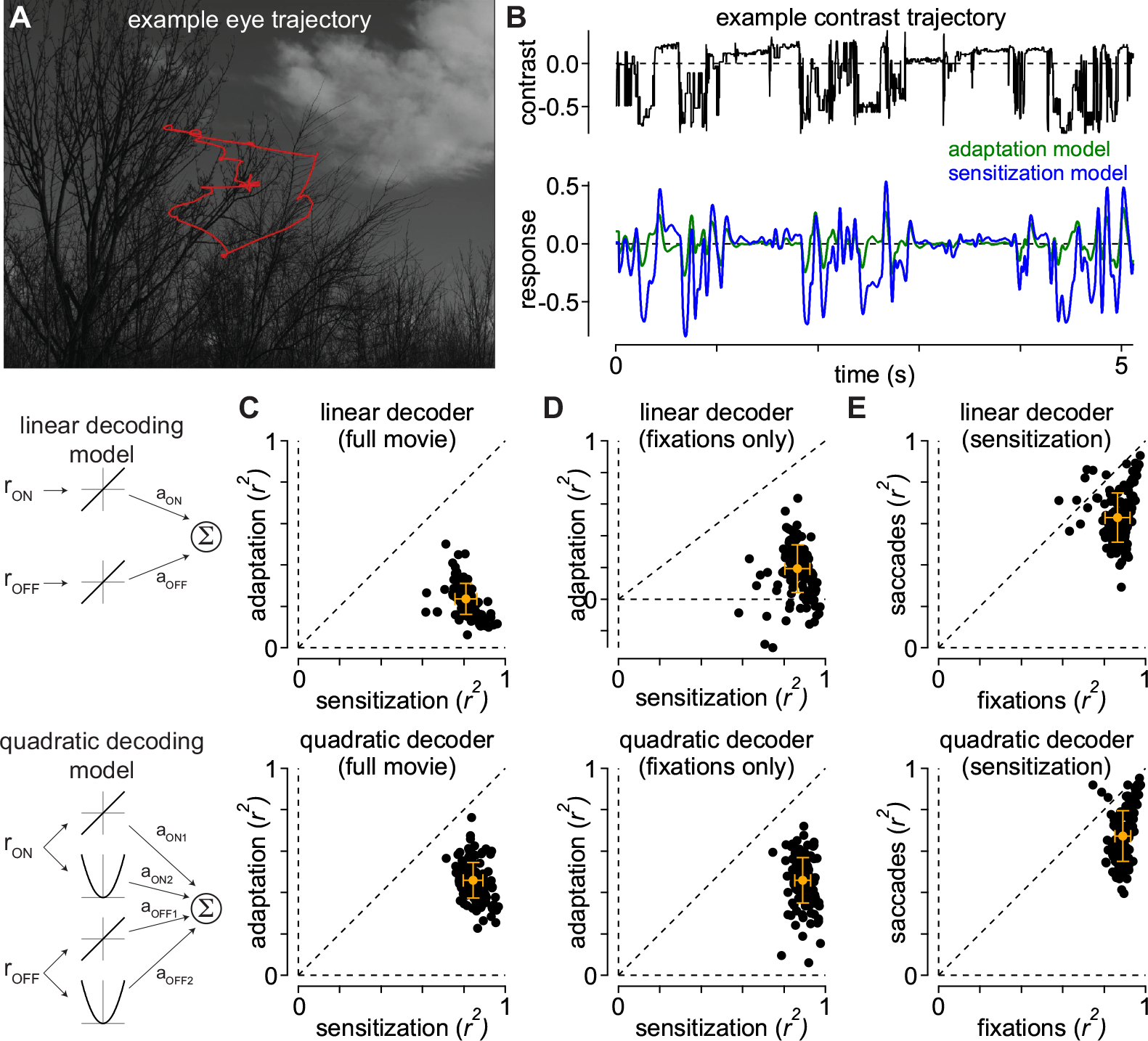
Sensitization increases the fidelity of encoding natural movies. (A) Example image from the DOVES database. The observer’s eye trajectory is shown in red. (B) *Top*, temporal contrast sequence from the eye movement data in (A). *Bottom*, responses of the adaptation and sensitization models to the example contrast sequence. (C) Performance of the sensitization (*x*-axis) and adaptation (*y*-axis) models at reconstructing 161 natural movies in the database. Performance was measured as the Pearson correlation between the stimulus and model predictions after adjusting for temporal lag. Performance for each movie is indicated by a black dot. Gray dot and bars indicate mean ± SD. The sensitization model outperformed the adaptation model for each of the movies. (D) Model performances as in (C), but restricted to periods of fixation. The sensitization model outperformed the adaptation model in each case. (E) Sensitization model performance for periods of fixation versus periods of eye motion. Predictive performance of the model was typically higher during periods of fixation.

The sensitizing model showed increased encoding accuracy for periods of fixation relative to periods of ballistic eye movements (movement *r*^2^, 0.63 ± 0.12; p = 2.9 × 10^−35;^ Figure 7E). This finding suggested that sensitization could play a particularly important role in vision during periods of fixation following the offset of global motion. We, thus, sought to determine whether background motion could evoke contrast sensitization with direct recordings from midget ganglion cells.

### Background motion evokes contrast sensitization in midget cells

To determine whether background motion elicited sensitization, we measured contrast responses in midget cells following the offset of a full-field moving texture (speed, 2 5-11 degrees s^−1^; duration, 1 s). The goal was to simulate, as closely as possible, the brief periods of fixation following eye movements and to test sensitivity during these fixation periods. We interleaved these recordings with measurements when the texture was stationary throughout the trial. The moving textures elicited an increase in spiking and a leftward shift in the contrast-response functions relative to the control condition in which the texture was stationary (Figure 8). On average, the shift was −25% contrast for spike recordings (−25.4 ± 4.4% contrast; n = 10 cells; p = 2.4 × 10^−2^) and −12% contrast for excitatory current recordings (−12.5 ± 5.1% contrast; n = 4 cells; p = 2.4 × 10^−2^).

**Figure 8.**
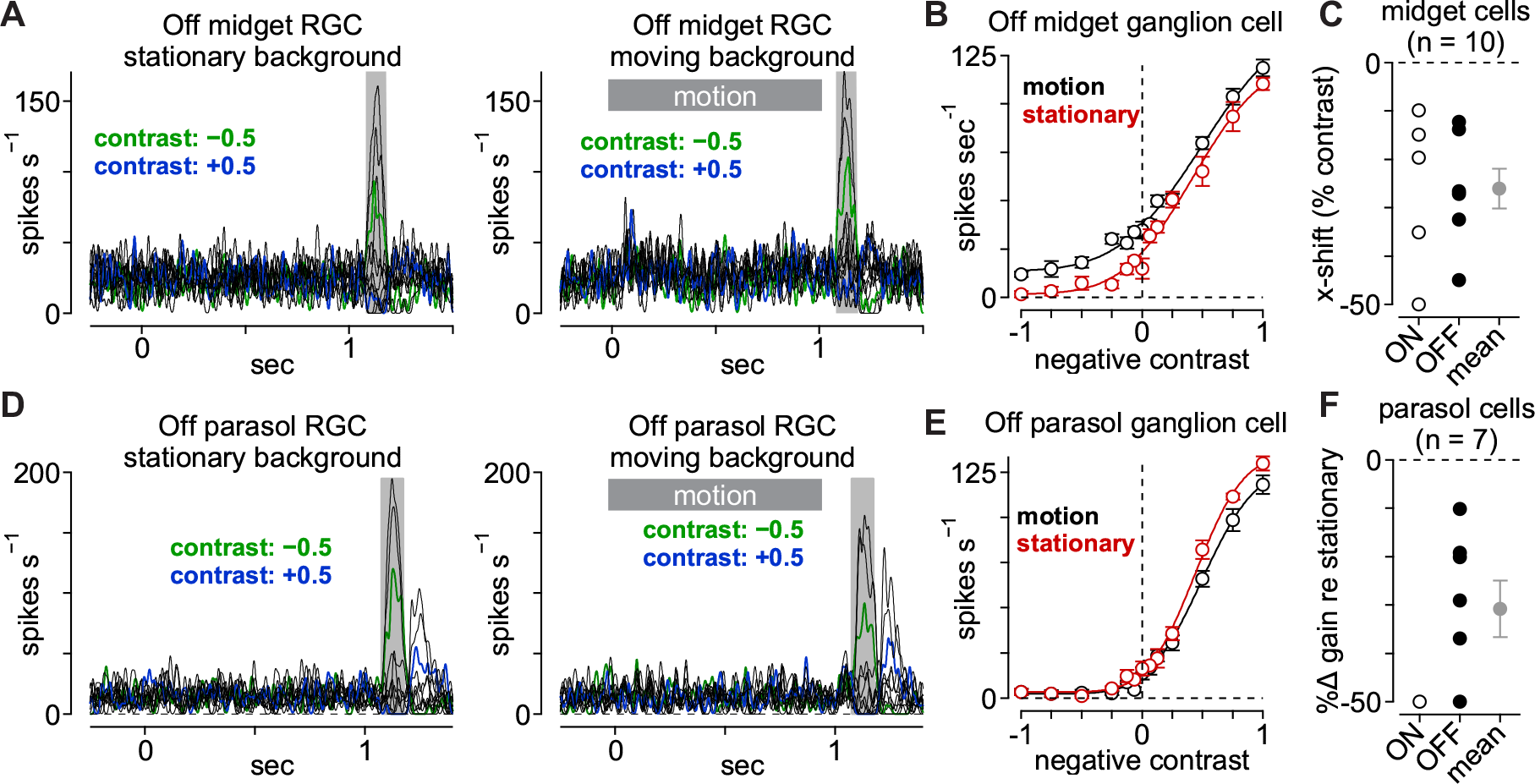
Background motion evokes contrast sensitization in midget cells. (A) Spike responses from an Off midget ganglion cell to a series of spots centered over the receptive field. Spots were either presented alone (left) or 50 ms following the offset of background motion (speed, 11 degrees s^−1^). Shaded regions indicate sampling windows. (B) Average spike rate across the shaded regions indicated in (A). The wide-field adaptation evoked a leftward shift in the contrast-response curve (black) relative to the unadapted control condition (red). (C) Horizontal shift (*x*-shift) in contrast-response function following background motion relative to control condition in which the background was stationary. Data are shown for On and Off midget ganglion cells (n = 10). Gray circle and bars indicate mean ± SEM. (D) Same as (A) for an Off parasol ganglion cell. (E) Same as (B) for the Off parasol cell in (D). The cell showed a decrease in spike output following the offset of background motion—the opposite pattern to that observed in the Off midget cell. (F) Change in gain in the contrast-response function following background motion relative to the control condition. On average, background motion elicited a decrease in gain of ~30% relative to the control condition in which the background was stationary (n = 7 cells). Gray circle and bars indicate mean ± SEM.

These data were consistent with our circuit model of contrast sensitization. The amacrine cell providing presynaptic inhibition to the midget bipolar cell adapted during background motion; at the offset of motion, the cell hyperpolarized and reduced presynaptic inhibition to the bipolar terminal. Thus, similar to circuits described in other vertebrates, the midget pathway could utilize presynaptic inhibition to account for self-motion (Olveczky et al., 2003; Baccus et al., 2008; Kastner and Baccus, 2013).

## DISCUSSION

Our results support a novel role for neural sensitization in primates relative to the function proposed in other species. Sensitizing cells are commonly thought to counteract the loss of responsiveness experienced by adapting cells during transitions from high to low variance environments (Kastner and Baccus, 2011). This hypothesis requires that sensitizing cells have an adapting counterpart that encodes similar information about the environment. Midget (parvocellular-projecting) ganglion cells are well known for their roles in both chromatic and achromatic vision (Crook et al., 2011; De Monasterio and Gouras, 1975; Derrington et al., 1984). Functional parallelism in the midget pathway is achieved by splitting signals between different classes of cone photoreceptor (L versus M) or bipolar cell (On versus Off) inputs to the midget cell receptive-field. Further, we found that both Onand Off-type midget cells exhibited sensitization (Figure 1-4, 8), and the primate retina lacks an adapting functional counterpart to midget cells with similar chromatic opponency or spatial acuity (Wässle, 2004); thus, sensitization does not counterbalance adaptation in another functionally parallel pathway.

Instead, our findings indicate that sensitization maintains the responsiveness of the midget pathway during dynamic visual processes, such as head or eye movements, that cause rapid fluctuations in light intensity on the retina. We base this conclusion on several key observations. First, sensitization was strongest following wide-field stimulation (Figure 1–4) or background motion (Figure 8). Second, sensitization persisted for >0.2 s (Figure 3), a period that roughly corresponds to the durations of fixations following eye movements in primates (reviewed in (Rayner, 1998)). Finally, sensitization greatly improved the fidelity of encoding natural movies, particularly during periods of fixation following ballistic eye motion (Figure 7). Thus, sensitization appears to play a unique and crucial role in neural coding in primates.

A parallel study also found evidence supporting the link between the sensitization mechanisms that we observed in midget ganglion cells and visual perception in humans (Naecker and Baccus, 2018). Subjects showed a significant enhancement in contrast sensitivity following the offset of wide-field motion; and this increase in sensitivity was manifest as a leftward horizontal shift in the perceptual input-output relationship, just as we observed in midget cells (compare Figure 2 in our study with Figure 5 of (Naecker and Baccus, 2018)). Together, these findings provide a rare example of a behavior that can be directly tied to a specific neural circuit motif.

### Distinct functions of adaptation and sensitization in primate retina

Our findings also speak to the roles of neural adaptation in the parasol and broad thorny ganglion cell pathways. Previous work proposed that adapting cells could produce a nearly optimal faithful encoding of sensory inputs (Fairhall et al., 2001). Our computational model, however, indicates that sensitizing circuits outperform adapting circuits in encoding natural movies (Figure 7). The improved reconstruction accuracy of the sensitizing model was consistent with a recent theoretical report indicating that sensitizing cells are better for encoding faithful representations of sensory input than adapting cells (Młynarski and Hermundstad, 2018). According to this paradigm, sensitizing cells such as midget ganglion cells would be useful for directly encoding information about the properties of the input (e.g., contrast, color). Adapting cells, on the other hand, are optimized for performing inference tasks (Wark et al., 2009; Młynarski and Hermundstad, 2018).

Adapting cells dynamically adjust their input-output properties to align with the recent stimulus distribution (Baccus and Meister, 2002; Smirnakis et al., 1997). These adjustments make the cells exquisitely sensitive to changes in stimulus statistics, allowing them to infer when salient properties of the environment change. For example, quickly detecting object motion is an ethologically relevant and phylogenetically ancient neural computation (Frost et al., 1990; Lettvin et al., 1959); by decreasing their responsiveness during periods in which the background is either stationary or coherently moving, adapting neural circuits would be poised to report when an object moves relative to the background (Olveczky et al., 2003; Puller et al., 2015). Interestingly, both adapting parasol and broad thorny ganglion cells have been implicated in motion processing (Manookin et al., 2018; Puller et al., 2015) and project to retinorecipient brain regions in the lateral geniculate body, superior colliculus, and inferior pulvinar that contribute significantly to motion vision (Rodieck and Watanabe, 1993; Crook et al., 2008; Kwan et al., 2018).

### Relationship to psychophysical measurements in humans

It has long been recognized that eye movements play important computational roles in visual processing (reviewed in (Martinez-Conde et al., 2004; Rucci and Victor, 2015)). Periods in which an image is stabilized on the retina cause that image to fade from perception (Troxler, 1804) and small fixational eye movements appear to counteract this fading (Rucci et al., 2007; Schütz et al., 2008). These eye movements can, how ever, produce large temporal fluctuations in contrast, particularly when viewing high contrast objects. This would, in turn, produce fading phenomena in cells that strongly adapt, such as parasol ganglion cells—a prediction that was confirmed with our computational model (Figure 7).

Neural mechanisms such as sensitization may serve to counteract adaptation by maintaining the sensitivity of certain visual pathways during eye movements. Indeed, our computational model and direct measurements indicated that contrast sensitization in the midget ganglion cell pathway was engaged well by background motion such as that observed during eye movements (Figure 7, 8). Thus, contrast sensitization might act to maintain sensitivity of image-forming visual pathways following eye movements that are commonplace in primate vision. Indeed, psychophysical studies in humans indicated that contrast sensitivity increases following both ballistic (saccade) and fixational eye movements (Rucci et al., 2007; Schütz et al., 2008). Moreover, this increase in sensitivity was limited to chromatic stimuli and high-spatial-frequency achromatic stimuli, mirroring our results in midget ganglion cells.

## METHODS

Experiments were performed in an *in vitro*, pigment-epithelium attached preparation of the macaque monkey retina (Manookin et al., 2015). Eyes were dissected from terminally anesthetized macaque monkeys of either sex (Macaca *fascicularis*, *mulatta*, and *nemestrina*) obtained through the Tissue Distribution Program of the National Primate Research Center at the University of Washington. All procedures were approved by the University of Washington Institutional Animal Care and Use Committee.

### Tissue Preparation and Electrophysiology

The retina was continuously superfused with warmed (32-35 °C) Ames’ medium (Sigma) at ~6-8 mL min^−1^. Recordings were performed from macular, mid-peripheral, or peripheral retina (2-8 mm, 10-30° foveal eccentricity), but special emphasis was placed on recording from more centrally located cells. Physiological data were acquired at 10 kHz using a Multiclamp 700B amplifier (Molecular Devices), Bessel filtered at 3 kHz (900 CT, Frequency Devices), digitized using an ITC-18 analog-digital board (HEKA Instruments), and acquired using the Symphony acquisition software package developed in Fred Rieke’s laboratory (http://symphony-das.github.io).

Recordings were performed using borosilicate glass pipettes containing Ames medium for extracellular spike recording or, for whole-cell recording, a cesium-based internal solution containing (in mM): 105 CsCH_3_SO_3_, 10 TEA-Cl, 20 HEPES,10 EGTA, 2 QX-314, 5 Mg-ATP, and 0.5 Tris-GTP, pH ~7.3 with CsOH, ~280 mOsm. Series resistance (~3-9 MΩ) was compensated online by 50%. The membrane potential was corrected offline for the approximately −11 mV liquid junction potential between the intracellular solution and the extracellular medium. Excitatory and inhibitory synaptic currents were isolated by holding midget ganglion cells at the reversal potentials for inhibitory/chloride (E_Cl_, ~−70 mV) and excitatory currents (E_cation_, 0 mV), respectively.

### Visual Stimuli and Data Analysis

Visual stimuli were generated using the Stage software package developed in the Rieke lab (http://stage-vss.github.io) and displayed on a digital light projector (Lightcrafter 4500; Texas Instruments) modified with custom LEDs with peak wavelengths of 405, 505 (or 475), and 640 nm. Stimuli were focused on the photoreceptor outer segments through a 10X microscope objective. Mean light levels were in the low to medium photopic regimes (~3 × 10^3^ − 3.4 × 10^4^ photoisomerizations [R*] cone^−1^ sec^−1^). Contrast values for contrast-response flashes are given in Weber contrast and for periodic stimuli in Michaelson contrast. All responses were analyzed in MATLAB (R2018a+, Mathworks).

For extracellular recordings, currents were wavelet filtered to remove slow drift and amplify spikes relative to the noise (Wiltschko et al., 2008) and spikes were detected using either a custom k-means clustering algorithm or by choosing a manual threshold. Whole-cell recordings were leak subtracted and responses were measured relative to the median membrane currents immediately preceding stimulus onset (0.25-0.5 s window). Summary data are presented in terms of conductance (*g*), which is the ratio of the current response (*I*) to the driving force:

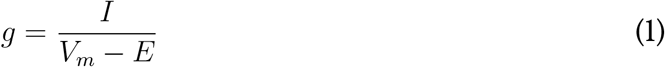

where *V*_*m*_ is the holding potential (in mV) and *E* is the reversal potential (in mV). Reversal potentials of 0 mV and −70 mV were used for excitatory and inhibitory inputs, respectively.

### Sensitization and adaptation models

We modeled spatiotemporal integration in bipolar cells and amacrine cells as the product of a Gaussian spatial filter and a biphasic temporal filter which was then passed through an input-output nonlinearity. The output of this nonlinear stage of the amacrine cell model was then passed through an adaptation stage; adaptation in the amacrine cell provided inhibitory input to the bipolar cell model prior to the output nonlinearity (Figure 6A). Following the subunit output, model midget ganglion cells and amacrine cells pooled (summed) inputs from bipolar cell subunits and the weights of these inputs were normalized by the subunit location relative to the receptive field center using a Gaussian weighting.

To estimate the excitatory and inhibitory circuit components for the computational model, we recorded excitatory and inhibitory synaptic currents from midget ganglion cells in response to a full-field Gaussian flicker stimulus. The contrast of each frame was drawn randomly from a Gaussian distribution and that value was multiplied by the average contrast. Average contrast was updated every 0.5 s and drawn from a uniform distribution (0.05-0.35 RMS contrast). The linear temporal filters (*F*) were calculated by cross-correlating the stimulus sequence (*S*) and the leak-subtracted response (*R*) (Baccus and Meister, 2002).

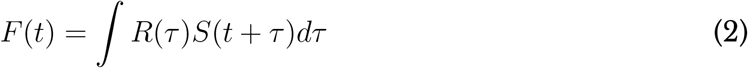

where *τ* is the temporal lag. These filters were then modeled as a damped oscillator with an S-shaped onset (Schnapf et al., 1990; Angueyra and Rieke, 2013):

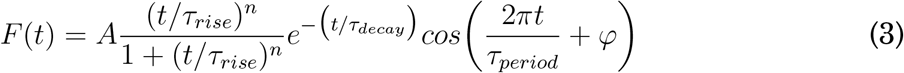

where *A* is a scaling factor, *τ*_*rise*_ is the rising-phase time constant, *τ*_*decay*_ is the damping time constant, *τ*_*period*_ is the oscillator period, and *φ* is the phase (in degrees).

The input-output nonlinearity was calculated by convolving the temporal filter (*F*) and stimulus (*S*) to generate the linear prediction (*P*).

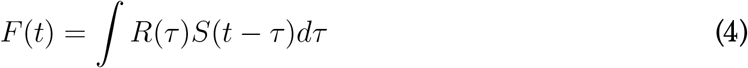

The prediction (*x*-axis) and response (*y*-axis) were modeled as a cumulative Gaussian distribution (Chichilnisky, 2001).

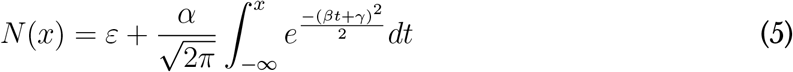

where *α* indicates the maximal output value, *ϵ* is the vertical offset, *β* is the sensitivity of the output to the generator signal (input), and *γ* is the maintained input to the cell.

The spatial component of the bipolar and amacrine cell receptive fields was modeled as a Gaussian function with a 2-SD width of 18 μm and 90 μm, respectively. Each midget ganglion cell was modeled as receiving input from a single bipolar cell, as is typically the case in the central retina. Sensitization parameters were determined by fitting linear-nonlinear model predictions relative to the excitatory currents recorded to the Gaussian flicker stimulus.

The amacrine cell providing direct inhibition to the midget ganglion cells is likely distinct from the cell providing presynaptic inhibition at the level of the midget bipolar cell (see Figure 5). Thus, our inhibitory synaptic recordings likely did not grant us direct access to the properties of the amacrine cell responsible for contrast sensitization. These recordings do, however, provide an estimate of the time-course of signals passing through the presynaptic amacrine cell to midget bipolar cells. Signals passing through this amacrine cell proceed from cone photoreceptors to bipolar cells and then to the amacrine cell in question before providing input to the midget bipolar cell. In the same way, the amacrine cell providing direct inhibition to midget ganglion cells must pass through an extra synapse. Thus, our recordings of direct synaptic inhibition were useful in approximating the time course of presynaptic inhibition at the midget bipolar terminal.

### Evaluating model performance to naturalistic movies

We evaluated the performance of the adaptation and sensitization models in reconstructing the naturalistic movie sequences using linear and quadratic decoding paradigms. To estimate stimulus contrast, the linear decoder (*f*_*LINEAR*_) summed the scaled outputs of the model On and Off midget ganglion cells:

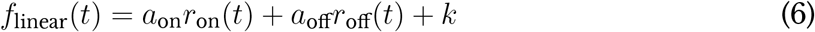

where *a*_*ON*_ and *a*_*OFF*_ are scaling constants and *k* is an offset constant. The quadratic model was similar in structure except that the response from each pathways was squared prior to summation:

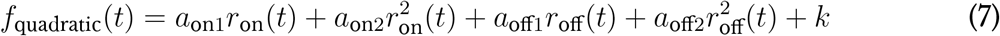

For each of the 161 movies in the database, the input stimulus was shifted to the peak of the midget temporal filter (~35 ms) and then scaling and offset coefficients were determined using least-squares curve fitting. The Pearson correlation was then calculated between the temporal trajectories of the model and the movie.

## ACKNOWLEDGEMENTS

We thank Shellee Cunnington, Mark Cafaro, and Jim Kuchenbecker for technical assistance. Tissue was provided by the Tissue Distribution Program at the Washington National Primate Research Center (WaNPRC; supported through NIH grant P51 OD-010425), and we thank the WaNPRC staff, particularly Chris English, for making these experiments possible. Fred Rieke, Raunak Sinha, Max Turner, and Will Grimes assisted in tissue preparation. We thank Jay Neitz and Fred Rieke for helpful discussions. We also thank Alison Weber and Jon Demb for feedback on a previous version of this manuscript. This work was supported in part by grants from the NIH (NEI R01-EY027323 to M.B.M.; NEI P30-EY001730 to the Vision Core), Research to Prevent Blindness Unrestricted Grant (to the University of Washington Department of Ophthalmology), Latham Vision Research Innovation Award (to M.B.M.), and the Alcon Young Investigator Award (to M.B.M.).

## AUTHOR CONTRIBUTIONS

Conceptualization, M.B.M.; Methodology, M.B.M.; Software, M.B.M.; Formal Analysis, M.B.M.; Investigation, M.B.M., T.R.A; Resources, M.B.M.; Data Curation, M.B.M., T.R.A.; Writing − Original Draft, M.B.M.; Writing − Review & Editing, M.B.M., T.R.A.; Visualization, M.B.M.; Supervision, M.B.M.; Project Administration, M.B.M.; Funding Acquisition, M.B.M. The ORCID number for M.B.M. is 0000-0001-8116-7619.

